# It takes two to tango: M-current swings with the persistent sodium current to set the speed of locomotion

**DOI:** 10.1101/2020.04.24.059311

**Authors:** Jeremy Verneuil, Cécile Brocard, Laurent Villard, Julie Peyronnet-Roux, Frédéric Brocard

## Abstract

The central pattern generator (CPG) for locomotion is set of pacemaker neurons endowed with inherent bursting driven by the persistent sodium current (*I*_NaP_). How they proceed to regulate the locomotor rhythm remained unknown. Here, in neonatal rodents, we identified a persistent potassium current, critical in regulating pacemakers and locomotion speed. This current recapitulates features of the M-current (*I*_M_); a subthreshold non-inactivating outward current blocked by XE991 and enhanced by ICA73. Immunostaining and mutant mice highlight an important role of axonal Kv7.2 channels in mediating *I*_M_. Pharmacological modulation of *I*_M_ regulates the emergence and the frequency regime of both pacemaker and CPG activities, and controls the speed of locomotion. Computational models captured these results and show howed an interplay between *I*_M_ and *I*_NaP_ that endows the locomotor CPG with rhythmogenic properties. Overall, this study provides fundamental insights into how *I*_M_ and *I*_NaP_ work in tandem to set the speed of locomotion.

## Introduction

Locomotion requires a recurrent activation of muscles with variable rhythm to adapt speed of movements as circumstances demand. In mammals, rhythmicity appears to be ensured by a network mainly localized in the ventromedial grey matter of upper lumbar segments [1, 2]. The rhythm-generating network is set of pacemaker cells endowed with intrinsic bursting activity in a frequency range similar to stepping rhythms [3, 4]. In exploring the ionic basis for rhythmogenesis, we identified the persistent sodium current (*I*_NaP_) as a critical current in burst-generating mechanism [5-7]. The immediate assumption was that the locomotor rhythm may emerge from neurons incorporating *I*_NaP_ as a “pacemaker” current. In line with this concept, inhibition of *I*_NaP_ abolishes locomotor-like activity in rodents [3, 8-10], salamanders [11] and disrupts locomotion in zebrafish [7, 12] or *Xenopus laevis* tadpoles [13]. Altogether, a picture emerges that the locomotor rhythm arises from a dynamic interplay between circuit-based activity and pacemaker burst-generating mechanisms with a critical role of *I*_NaP_ for the initiation of bursts [4, 9].

Among general principles of rhythmogenesis, outward conductances are required to repolarize bursts [14]. In vertebrates, the Ca^2+^-activated K^+^ current (*I*_*KCa*_) appears important in repolarizing bursts to regulate the locomotor rhythm [15-18], among other hyperpolarizing currents mediated by Kv1.2 [19] and A-type K^+^ channels [20] or Na^+^/K^+^ pumps [21-23]. Considering *I*_NaP_ as a subthreshold persistent conductance critically engaged in the burst initiation, it might be efficiently counteracted by a subthreshold persistent potassium current to end bursts of pacemakers and thereby to regulate the locomotor cycle. The Kv7 (KCNQ) channels mediate a unique subthreshold persistent outward current named *I*_M_ [24]. The Kv7 channels are widely expressed in the CNS and several pieces of evidence reveal the existence of *I*_M_ in spinal motoneurons, presumably mediated by heteromeric Kv7.2/Kv7.3 channels [25-29]. However, their expression at the level of the locomotor central pattern generator (CPG) and their functional role in locomotion remain unexplored.

The present study characterizes the functional expression of *I*_M_ in locomotor-related interneurons and identifies Kv7.2 channels as its molecular constituent. We show that the modulation of *I*_M_ adjusts *I*_NaP_ amplitude, regulates the emergence of pacemaker properties, their burst repolarization, the frequency regime of the locomotor CPG and the speed of locomotion. In sum, we provide the first description of *I*_M_ in a vertebrate motor CPG and describe its dynamic interplay with *I*_NaP_ as a fundamental mechanism in shaping bursting activity of neurons and networks that control rhythmic locomotor output.

## Results

### M-current regulates the speed of locomotion

To study the role of *I*_M_ in the locomotor behavior of juvenile rats (15-to 21-day-old), potent and selective Kv7 channel modulators were injected intraperitoneally at a concentration commonly used for a systemic modulation of *I*_M_ [30-32]. The regular pattern of walking was captured with the CatWalk system before and after 30 min of drug administration (**Fig 1**). Control experiments with DMSO discarded potential effects of the vehicle (*P* > 0.05, in grey, **Fig 1A–1C**). Rats walked faster when the broad-spectrum activation of Kv7.2**–**Kv7.5 channels occurred by injecting retigabine [(5 mg/kg see [33]], which was related to a shorter stance phase (*P* < 0.05, in green, **Fig 1A–1C**). The broad-spectrum inhibition of Kv7 channels by linopirdine (3 mg/kg) [34] had opposite effects; i.e. rats walked slower, attributable to a longer stance phase (*P* < 0.05, in yellow, **Fig 1A–1C**). A second set of experiments focused on the role of Kv7.2/3 subunits by means of more selective drugs. The potent Kv7.2/3 channel opener ICA73 (5 mg/kg) [35] (*P* < 0.05, in blue, **Fig 1A–1C**) and the Kv7.2/3 channel blocker XE-991 (5 mg/kg) [34] (*P* < 0.05, in red, **Fig 1A–1C**) reproduced retigabine and linopirdine effects, respectively. Noteworthy, none of the Kv7 modulators affected the inter-limb coordination or gait (*P* > 0.05; **S1 Fig**). Altogether, these results support the concept that *I*_M_, presumably mediated by Kv7.2 and/or Kv7.3 channels, is an important and physiologically relevant regulator of the “clock” function of the locomotor network without interfering with the dynamic postural control.

**Figure 1:**
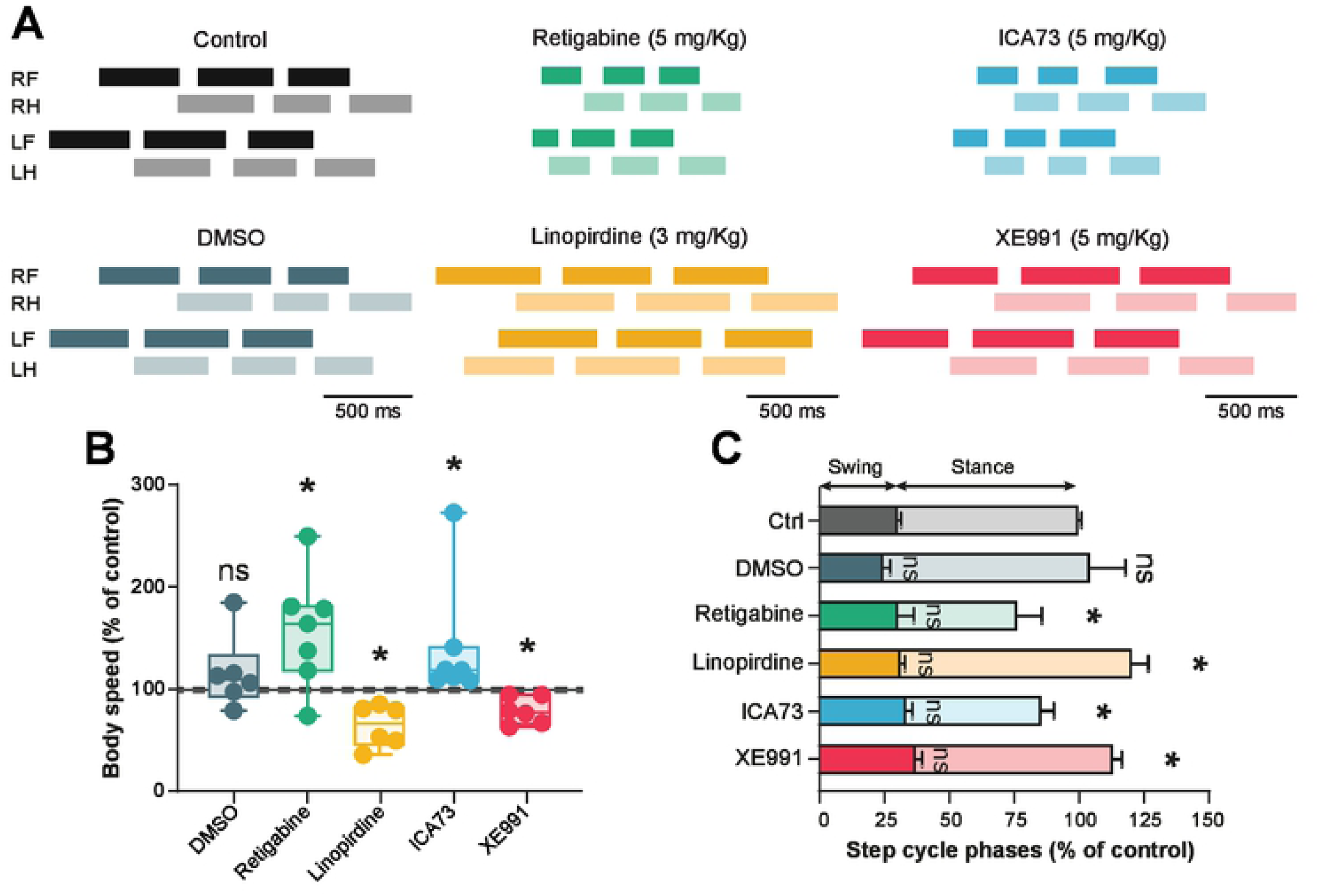
*I*_M_ sets the speed of locomotion. (**A**) Representative footfall diagrams during CatWalk locomotion of juvenile rats (15-to 21-d-old) before (*black*) and 30 min after acute i.p. administration of DMSO (*grey, n* = 6 rats), retigabine (5 mg/kg, *green, n* = 7 rats), linopirdine (3 mg/kg, *yellow, n* = 6 rats), ICA73 (5 mg/kg, *blue, n* = 7 rats) or XE991 (5 mg/kg, *red, n* = 6 rats). (RF, right forelimb; RH, right hindlimb; LF, left forelimb; LH, left hindlimb). The stance phase is indicated by horizontal bars and the swing phase by open spaces. (**B**) Normalized changes of the body speed induced by the above-mentioned drugs. *G*rey shading indicates the 95% confidence intervals of control values. \**P* < 0.05, comparing animals before and after drug administration; Wilcoxon paired test. (**C**) Normalized changes of swing and stance phases expressed as a percentage of the total step cycle. \**P* < 0.05, comparing data collected before and after drug administration; Wilcoxon paired test. Data in **C** are mean ± SEM. Underlying numerical values can be found in the Figure 1 source data.

### Ubiquitous axonal expression of Kv7.2 channels in interneurons from the locomotor CPG region

Modulation of the step cycle through systemic administration of drugs may involve changes in the excitability of midbrain circuits setting the speed of locomotion [36, 37]. To investigate a putative modulation of *I*_M_ at the level of the spinal locomotor CPG, we studied the expression of Kv7.2/3 channels in ventromedial interneurons from upper lumbar segments (L1–L2), where components of the locomotor rhythm-generating network are located [1, 2]. Immunostaining substantiates the presence of Kv7.2 channels in all ventromedial interneurons, particularly in the distal part of axonal initial segments (AISs) identified by their expression of Nav channels (**Fig 2A–2C, 2G and 2H**). Similar Kv7.2-immunostaining profile was observed in lumbar motoneurons (**S2A–S2C, S2G and S2H Fig**). Conversely to Kv7.2, Kv7.3 subunit was virtually not detectable or weakly expressed in only a few ventromedial interneurons (**Fig 2D–2G**). On the other hand, Kv7.3-immunostaining distinctively labeled the distal part of AISs in almost half of lumbar motoneurons (**S2D–S2H Fig**). In sum, L1-L2 ventromedial interneurons ubiquitously express Kv7.2-containing channels, suggestive of the existence of *I*_M_ in the locomotor CPG.

**Figure 2:**
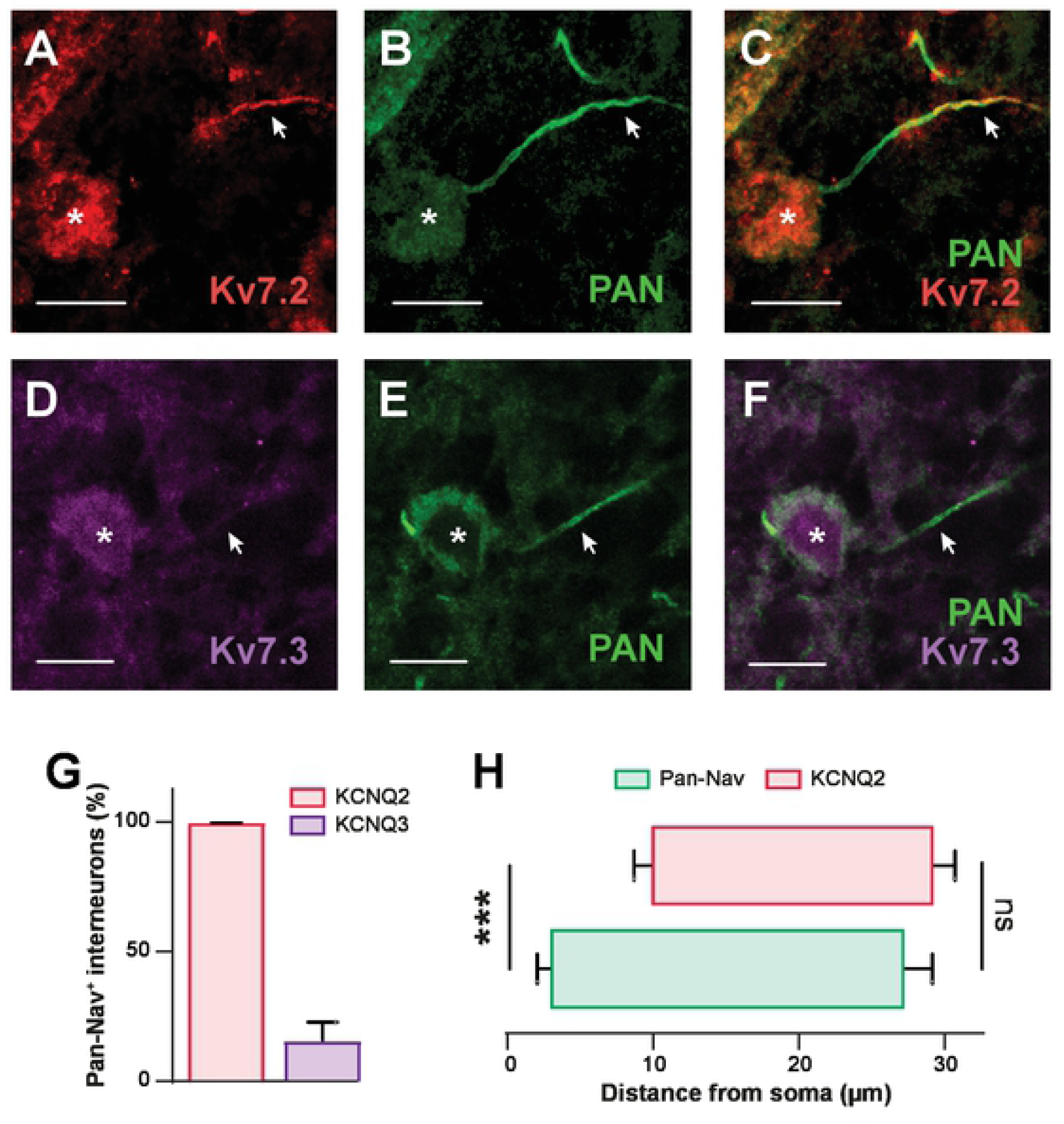
Lumbar interneurons within the locomotor CPG region express Kv7.2-containing channels. (**A-F**) Immunostaining of lumbar (L1-L2) ventromedial interneurons from juvenile rats (n = 3 rats) against Kv7.2 (**A**, n = 216 cells) or Kv7.3 (**D**, n = 170 cells) along the axon initial segment (AIS) labeled by the Pan-Nav antibody (**B**,**E**). Kv7.2 and Pan-Nav are merged in (**C**), and Kv7.3 and Pan-Nav are merged in (**F**). Asterisks indicate the nucleus position and arrowheads the AIS. Scale bars = 10 µm. (**G**) Group means quantification of the proportion of Pan-Nav positive interneurons expressing Kv7.2 or Kv7.3 channels. (**H**) Group means quantification of the start and end positions of Pan-Nav and Kv7.2 immunolabelings along the axonal process from the soma (n = 10 cells). ***P < 0.001, comparing start or end positions between groups; Mann-Whitney test. Data are mean ± SEM. Underlying numerical values can be found in the figure 2 source data.

### Characterization of *I*_M_ in the rhythm-generating kernel for locomotion

To assess whether interneurons from the CPG region exhibit a functional *I*_M_, we isolated *I*_M_ by whole-cell patch clamp recordings under TTX (1 µM). By using the standard relaxation protocol, we measured *I*_M_ as the amplitude of the tail inward current evoked by stepping down voltage from -10 mV (**Fig 3A**). All L1-L2 ventromedial neurons displayed an electrophysiological signature of *I*_M_. From the current-voltage (*I-V*) relationship fitted with a standard Boltzmann function (in black, **Fig 3B**), the threshold for activation (*V*_T_) was positive to -67.3 ± 1.9 mV, and its amplitude increased steeply (slope factor k: 4.4 ± 0.6) for larger voltage steps with a mid-point of activation (V_1/2_) at -43.6 ± 1.5 mV and then plateaued above -30 mV. The peak amplitude of *I*_M_ was in mean of 79.2 ± 7.4 pA. We further characterized *I*_M_ pharmacologically (**Fig 3C**). The *I*_M_-enhancer ICA73 (10 µM) increased the holding current, the magnitude of *I*_M_ and hyperpolarized its *V*_T_ and V_1/2_ activation (*P* < 0.05, in blue, **Fig 3B–3D**). The *I*_M_-enhancer retigabine (100 nM) replicated effects of ICA73 on *I*_M_ (**S3A–S3F Fig**). The *I*_M_-blocker XE991 (10 µM) markedly reduced the holding current and virtually abolished *I*_M_ (*P* < 0.05, in red, **Fig 3B–3D**). To confirm that CPG for locomotion express *I*_M_, we recorded from transgenic mice Hb9:GFP-positive ventromedial interneurons that are considered as components of the locomotor rhythm-generating circuit [38, 39]. All Hb9 cells displayed similar biophysical properties of *I*_M_ that was abolished by XE991 (**Fig 3E**). Given the strong expression of Kv7.2 in interneurons, we evaluated the contribution of Kv7.2 to *I*_M_ with knock-in mice bearing the T274M Kv7.2 mutation associated with Ohtahara syndrome [40]. Because this mutation is homozygous lethal we studied *I*_M_ on Kv7.2^Thr274Met/+^ animals. We found in heterozygous Kv7.2^Thr274Met/+^ mice that *I*_M_ was about halved in amplitude relative to Kv7.2^+/+^ littermates (**Fig 3F**). Altogether, these results support the expression of a non-inactivating K^+^ current corresponding to *I*_M_ in the locomotor CPG, presumably carried by Kv7.2-containing channels.

**Figure 3:**
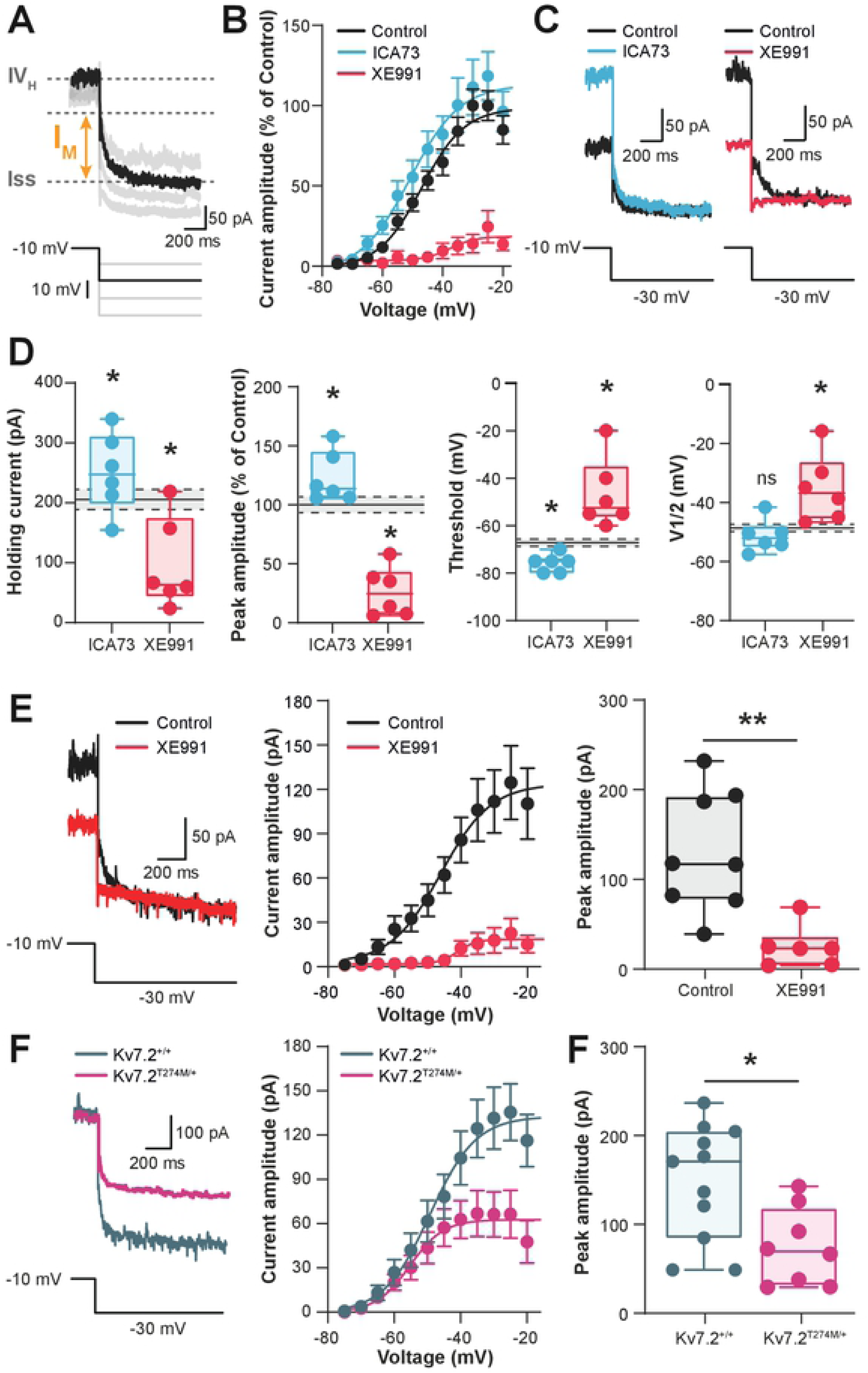
Characterization of *I*_M_ in interneurons of the locomotor CPG region. (**A**) Representative inward current relaxation induced by the standard *I*_M_ deactivation voltage protocol. The current relaxation (*I*_M_) was measured as the difference between the fast transient (*I*_*VH*_) and the steady-state current (*Iss*). (**B**) Boltzmann fitted current-voltage relationships of *I*_M_ normalized to control values and established before and after bath-applying XE991 (10 µM, red) or ICA73 (10 µM, blue). Error bars represent ± SEM. (**C**) Representative deactivation of *I*_M_ before and after adding the above-mentioned drugs. (**D**) Boxplots quantification of biophysical properties of *I*_M_. (n = 6 cells). Dashed lines with grey shading indicate the 95% confidence intervals of control values. \**P* < 0.05, \*\*\**P* < 0.001, comparing data collected before and after adding the above-mentioned drugs; Wilcoxon paired test. (**E**) Representative deactivation (*left*), Boltzmann fitted current-voltage relationships (*middle*) and amplitude (*right*) of *I*_M_ recorded in Hb9^+^ interneurons under control conditions (n = 8 cells, black) or in the presence of XE991 (10 µM, n = 6 cells, red). \**P* < 0.05; Mann-Whitney test. (**F**) Representative deactivation (*left*), Boltzmann fitted current-voltage relationships (*middle*) and amplitude (*right*) of *I*_M_ recorded in interneurons from wild-type (n = 11 cells, grey) and Kv7.2^Thr274Met/+^ mutant mice (n *=* 8 cells, purple). \**P* < 0.05; Mann-Whitney test. Underlying numerical values can be found in the figure 3 source data.

### M-current sets the neuronal excitability and gates pacemaker bursting mode

We characterized the role of *I*_M_ on membrane properties of L1-L2 ventromedial interneurons recorded from neonatal rats. As a first observation, the *I*_M_-enhancer ICA73 hyperpolarized cells (in blue, **S4A Fig**; routinely compensated to -60 mV) along with a drop of the input resistance (*P* < 0.05 **Table 1**). Therefore interneurons became less excitable (higher rheobase; *P* < 0.05, **Table 1**) and produced fewer spikes (*P* < 0.05, in blue, **Fig 4A**) without any changes in parameters of the action potential (*P* > 0.05, **Table 1**). The f-I curve was thus shifted to the right (**Fig 4B**). The reversibility of these effects when the *I*_M_-blocker XE991 was applied, emphasized the dependence of ICA73 on Kv7 channels (**S4A–S4D Fig**). Consistent with this, the Kv7-opener retigabine reproduced effects of ICA73 on electroresponsive properties (**S3G and S3H Fig and Table 1**). We further tested the functional implication of *I*_M_ by using the *I*_M_-blocker XE991 alone. The *I*_M_-blocker *per se* neither depolarised the resting membrane potential nor increased the neuronal excitability or the firing rate (*P* > 0.05, in red, **Fig 4A and 4B and Table 1**). Thus, the basal excitability was not affected by XE991. However, *I*_M_-blockers such as XE991 have the peculiarity to be voltage-dependent blockers with higher affinity at positive potentials and thereby are very poor inhibitors at perithreshold potentials [41]. Furthermore, the inhibition develops slowly upon depolarization within a timescale of minutes [42].

**Table 1:** Effects of drugs targeting Kv7 channels on passive and active membrane properties of interneurons. Values are means ± SEM; *n*, number of cells. Mean firing frequency was measured at twofold the rheobase. \**P* < 0.05, \*\*\**P* < 0.001, comparing data collected before and after bath-applying drugs mentioned above; Wilcoxon paired test.

|  | Vrest (mV) | Input Resistance (MΩ) | Rheobase (pA) | Spike threshold (mV) | Spike amplitude (mV) |
| --- | --- | --- | --- | --- | --- |
| Control | -57.8 ± 1.2 | 1147.3 ± 121.3 | 11.3 ± 1.4 | -45.0 ± 1.1 | 55.5 ± 2.2 |
| ICA73 (n=6) | -65.4 ± 2.8 * | 694.3 ± 222.4 * | 28.3 ± 10.7 * | -49.1 ± 1.4 (ns) | 48.1 ± 2.6 (ns) |
| Retigabine (n=9) | -70.7 ± 1.7 ** | 869.4 ± 103.2 * | 17.2 ± 3.8 * | -47.1 ± 1.9 (ns) | 48.4 ± 1.8 (ns) |
| XE991 (n=7) | -57.1 ± 1.8 (ns) | 979.1 ± 150.8 (ns) | 17.1 ± 2.6 (ns) | -43.7 ± 1.7 (ns) | 57.9 ± 4.0 (ns) |

**Figure 4:**
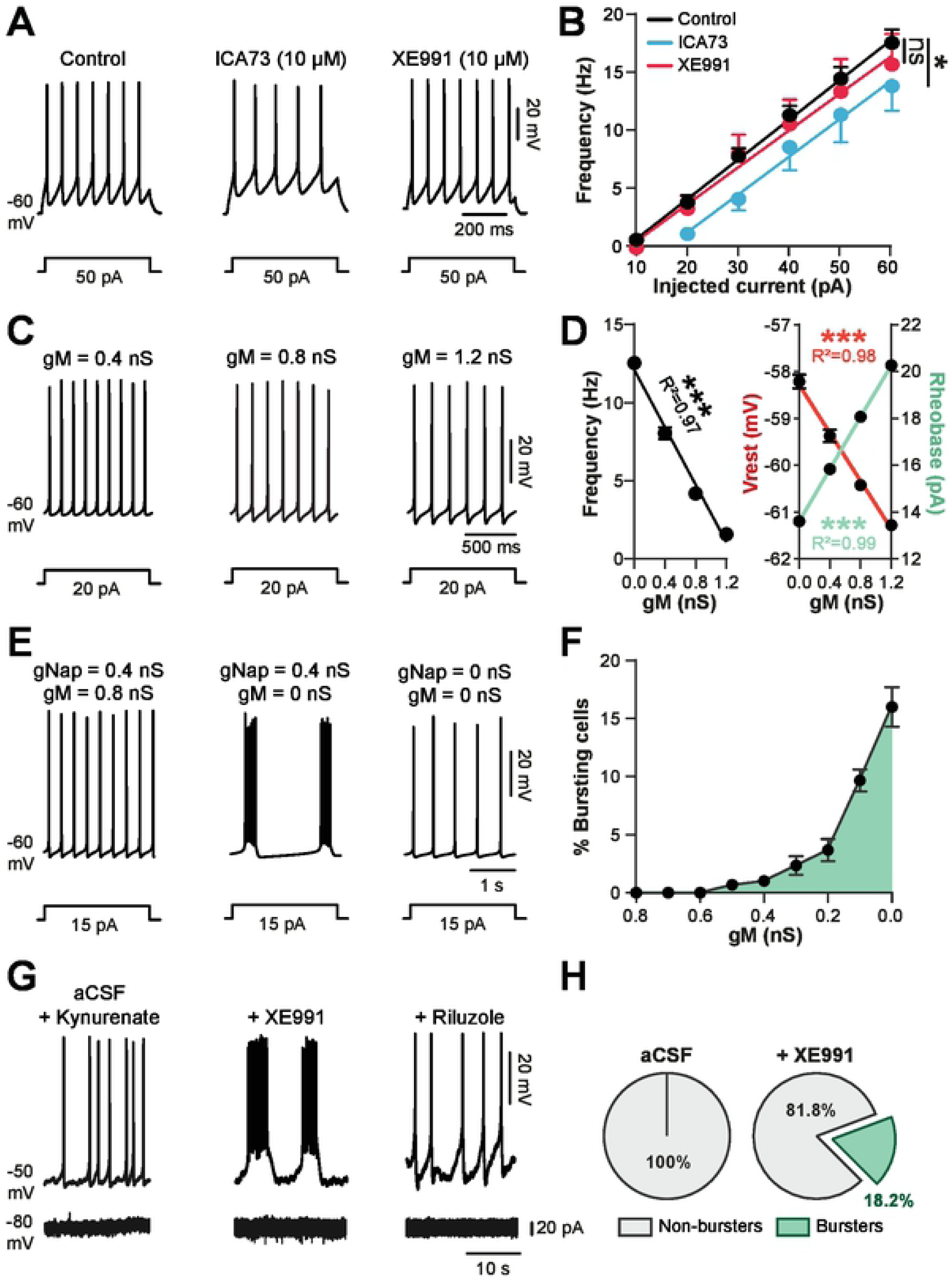
*I*_M_ regulates both the excitability and the firing pattern of L1-L2 ventromedial interneurons. (**A**,**B**) Spiking activity of ventromedial interneurons (L1-L2) to a near threshold depolarizing pulse (**A**) with its respective frequency-current relationships (**B**) before and after bath-applying ICA73 (10 µM, n = 7 cells, blue) or XE991 (10 µM, n = 7 cells, red). Continuous lines represent the best-fitting linear regression. \*\*\**P* < 0.001 comparison of the fits before and after adding the above-mentioned drugs. (**C**) Spiking activity from a single-neuron model at three different values of *g*_M_. The injected current pulses in **A** and **C** are indicated below voltage traces. (**D**) Mean plots of the firing rate, rheobase and resting membrane potential as function of *g*_M_ (in the heterogeneous population model of 50 neurons). The continuous line is the best-fitting linear regression. \*\*\**P* < 0.001, Spearman correlation test. r indicates the correlation index. (**E**) Typical switch of the firing pattern from spiking to bursting in a single-neuron model when *g*_M_ was switched “off”; simulated blockade of *I*_NaP_ (*g*_NaP_ = abolished bursting activity. (**F**) Dependence of the percentage of bursting cells on *g*_M_ in the heterogeneous population model of 50 neurons. (**G**) On top, representative voltage traces of a ventromedial interneuron (L1-L2) intracellularly recorded in the presence of kynurenate (1.5 mM) before (*left*) and after (*middle, red*) bath-applying XE991 (10 µM). A subsequent application of riluzole (5 µM) abolished bursts (*right*). At bottom, current traces of the cell illustrated and voltage clamped at - 80 mV. (**H**) Proportion of burster and non-burster interneurons before and after bath-applying XE991 (n = 22 cells). Data are mean ± SEM. Underlying numerical values can be found in the figure 4 source data.

To overcome this experimental limitation, we studied the theoretical effect of *I*_M_ by modeling *I*_M_ in a Hb9 cell model that we previously used [9]. The model was supplemented by *I*_M_ derived from our voltage-clamp recordings. A population of 50 uncoupled interneurons was simulated with a randomized normal distribution of neuronal parameters (see methods). Simulated neurons reproduced key features of the biological responses to stepwise depolarizing currents with firing rate in the range of our experimental data (**Fig 4C and 4D**). The increase of the M-conductance (*g*_M_) in the model qualitatively captured the modulation of neuronal excitability observed experimentally with *I*_M_-enhancers ICA73 or retigabine; the resting membrane potential hyperpolarized, the rheobase increased and firing rate decreased (*P* < 0.001, **Fig 4C and 4D**). However, in contrast to our electrophysiological recordings with the *I*_M_-enhancers XE991, decreasing *g*_M_ in the model predicted a depolarization of *V*_rest_, while the firing rate and the rheobase would go up and down, respectively (*P* < 0.001, **Fig 4C and 4D**). Another remarkable effect of modeling a decrease of *g*_M_ was the gradual transfer to a bursting mode in a small proportion of neurons to reach 17 % of bursters when *g*_M_ was switched off (**Fig 4E and 4F**). Bursts disappeared when *I*_NaP_ was zeroed (**Fig 4E**). To evaluate this computational prediction, we tested XE991 on cells intracellularly recorded and constantly depolarized with a suprathreshold current (**Fig 4G**). In this condition, XE991 caused a transition from tonic spiking to bursting in ∼18% of the interneurons recorded (**Fig 4G and 4H**). Bursts were abolished by the *I*_NaP_-blocker riluzole (**Fig 4G**). The insensitivity of bursts to kynurenic acid (1.5 mM; blocker of the fast glutamatergic transmission) or the lack of rhythmic currents in voltage clamp recordings, precluded a role of network inputs in the emergence of bursts (**Fig 4G**).

Overall, these data indicate an important role of *I*_M_ in setting the excitability and firing properties of interneurons within the locomotor CPG region notably by impeding the initiation of bursts mediated by *I*_NaP_.

### Interneurons balance *I*_M_ and *I*_NaP_ to trigger pacemaker bursting mode

The emergence of *I*_NaP_-dependent bursting cells when Kv7 channels are blocked suggests that *I*_M_ might counteract *I*_NaP_ to regulate pacemaker properties. To test this possibility, voltage-clamp recordings were performed to examine the degree of interaction between the two currents. In response to very slow voltage ramps, ventromedial interneurons from neonatal rats displayed a large inward current attributable to *I*_NaP_ [**Fig 5A**; see [5]]. In the presence of *I*_M_-blocker XE991, *I*_NaP_ was higher in amplitude while *V*_T_ and V_1/2_ did not change (*P* < 0.05, **Fig 5A and 5B**). These results show that biophysical properties of *I*_M_ are well suited to counteract the depolarizing drive furnished by *I*_NaP_. Here we speculate that the combination of *I*_M_ and *I*_NaP_ currents is a possible mechanism in controlling bursting dynamics. We evaluated this assumption, by varying the maximal conductance levels of *g*_M_ and *g*_NaP_ in the heterogeneous population model composed of 50 uncoupled Hb9-type interneurons (**Fig 5C**). None of the neurons exhibited bursting at base levels of *g*_M_ (0.8 nS) and *g*_NaP_ (0.4 nS). We found that a population bursting activity could be triggered by either a reduction of *g*_M_ or an increase of *g*_NaP_ in turn (**Fig 5C**). However, the striking observation was the synergistic effect of reducing *g*_M_ and increasing *g*_NaP_ conductances on the generation of bursts (**Fig 5D**). Compared to non-bursters, bursters were distinguished by a more negative V_1/2_ of *I*_NaP_ but displayed a similar V_1/2_ of *I*_M_ (*P* < 0.001, **Fig 5E**).

**Figure 5:**
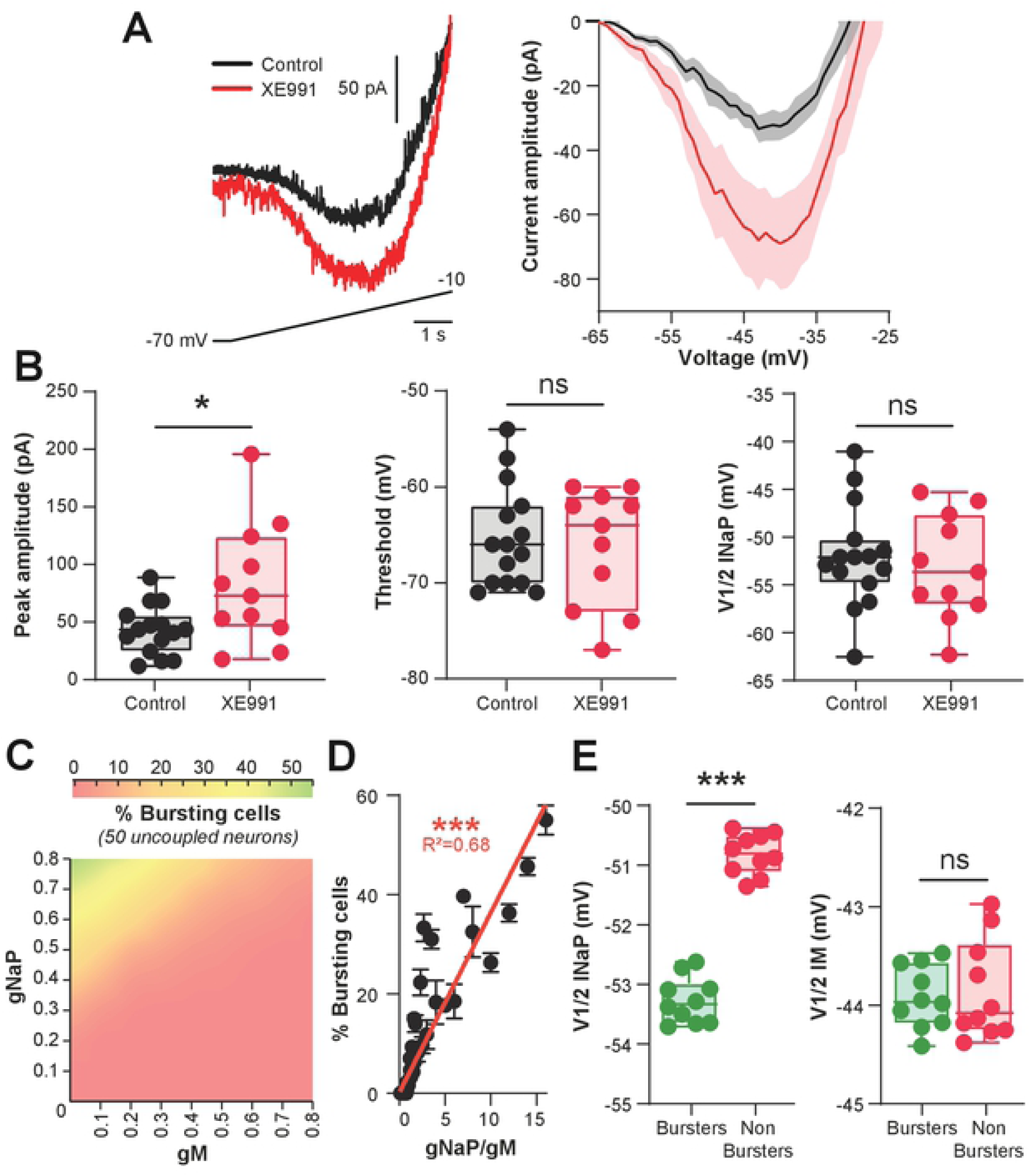
A balance of *I*_M_ and *I*_NaP_ gates bursting mode of CPG interneurons. ((**A**) On the left, raw traces of leak-subtracted persistent sodium current (*I*_NaP_) evoked in L1-L2 ventromedial interneuron by a slow voltage ramp, under control condition (black) and in the presence of XE991 (10 µM, red). On the right, mean leak-subtracted *I*_NaP_ recorded in neurons as a function of command voltage under control conditions (n = 15 cells, black) and in the presence of XE991 (n = 11 cells, red). The grey and red shadings indicate the 95% confidence interval. (**B**) Boxplots quantification of the biophysical properties of *I*_NaP_. \**P* < 0.05; Mann-Whitney test. (**C**) Dependence of the percentage of bursting pacemaker cells on *g*_M_ and *g*_NaP_ in the heterogeneous population model of 50 neurons. Results are pooled from 10 simulations. (**D**) Percentage of bursting cells as function of the *g*_NaP_ / *g*_M_ ratio. Each dot represents average values of 10 simulations for a fixed *g*_NaP_ / *g*_M_ ratio. Continuous line is the best-fitting linear regression. \*\*\**P* < 0.001, Spearman correlation test. r indicates the correlation index. (**E**) Boxplots quantification of V_1/2_ of *I*_NaP_ and *I*_M_ for burster and non-burster groups. Each dot represents one simulation in the heterogeneous population model of 50 neurons. Parameters values for *g*_NaP_ and *g*_M_ were set at 0.8 nS and 0.1 nS, respectively. \*\*\**P* < 0.001; Mann-Whitney test. Underlying numerical values can be found in the figure 5 source data.

Taken together, these results suggest that most of CPG interneurons are endowed with the intrinsic ability to switch from spiking to bursting behavior through a sliding balance between *I*_NaP_ and *I*_M_.

### M-current controls bursting dynamics of pacemaker cells

Our modeling study combined with electrophysiological data supports fine modulation of *g*_M_ as a key mechanism for the emergence of pacemaker cells. Here, we used the model as a tool to delineate the role of *I*_M_ in bursting dynamics. To study dependence of bursting characteristics on *I*_M_, we simulated a negative-voltage shift of *I*_NaP_ activation (V_1/2_ = -54 mV) to convert a tonic cell into burster as a result of reducing [Ca^2+^]_o_ [9]. As previously described, the burst period and the burst duration decreased as the neuron was depolarized (**Fig 6A**). When *g*_M_ was omitted, the burst duration as well as the interburst interval increased (*P* < 0.001, **Fig 6B and 6C**). Opposite effects on burst timing was observed when *g*_M_ was increased (*P* < 0.001, **Fig 6C**). Notably, at high level of *g*_M_, a subset of bursting cells in the heterogeneous population model switched to tonic firing (**S5A and S5B Fig**). To determine the dynamic contributions of *I*_M_ and *I*_NaP_ to the bursting activity, we examined changes of *g*_M_ and *g*_NaP_ at specific time points during the burst itself and during interburst intervals (**Fig 6D**). During the beginning of each burst, *g*_NaP_ preceded the activation of *g*_M_. Over the course of the burst, a slow decrease of *g*_NaP_ was observed whereas *g*_M_ slightly increased. At the end of the burst, *g*_M_ and *g*_NaP_ relaxed to a baseline level over the duration of an interburst interval. Altogether, these results suggest a scenario in which *I*_NaP_ initiates the burst and *I*_M_ contributes to the normal oscillatory activity of pacemakers by counteracting *I*_NaP_ during the burst.

**Figure 6:**
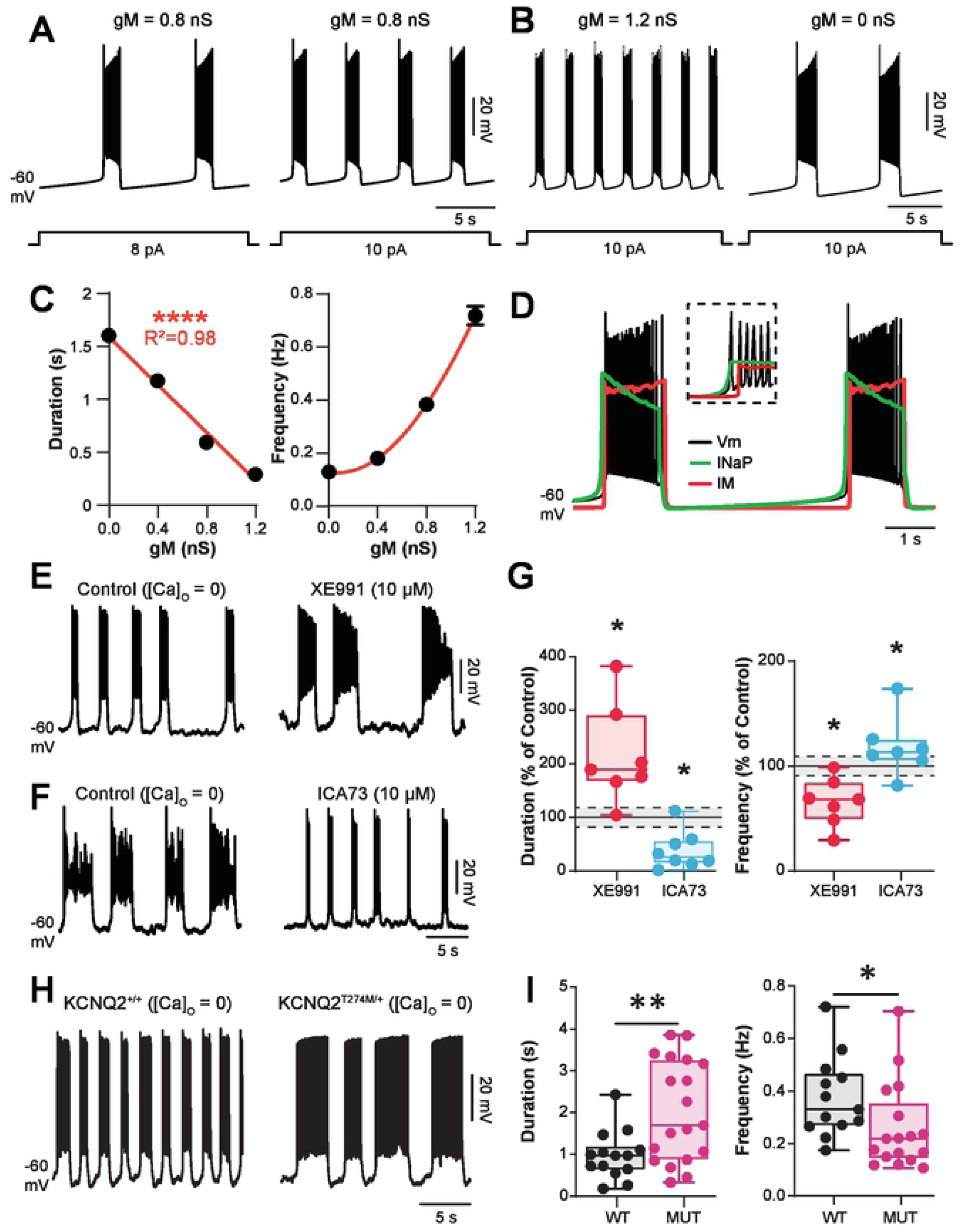
*I*_M_ controls burst dynamics. (**A**,**B**) Firing behavior from a bursting pacemaker neuron model in response to incrementing depolarizing current injections (**A**) or for two different values of *g*_M_ (**B**). The injected current pulses are indicated below voltage traces V_1/2_ *I*_NaP_ = -54mV. (**C**) Duration and frequency of bursts as function of *g*_M_ in the heterogeneous population model of 50 neurons. Results are pooled from 10 simulations. (**D**) Dynamics of the neuronal membrane potential (*black*), the *I*_NaP_ (*green*) and the *I*_M_ (*red*) during bursting activity of a single pacemaker neuron model. The broken-line box highlights dynamics of *I*_NaP_ and *I*_M_ at the onset of bursts. (**E**,**F**) free-[Ca^2+^]_o_ saline-induced bursting activity recorded intracellularly from L1-L2 ventromedial interneurons before and after XE991 (10 µM, n = 7 cells) (**E**) or ICA73 (10 µM, n = 7 cells) (**F**). (**G**) Normalized changes of burst parameters. *D*ashed lines with grey shading indicate the 95% confidence intervals of control values. \**P* < 0.05, comparing data before and after the above-mentioned drugs; Wilcoxon paired test. (**H**) Free-[Ca^2+^]_o_ saline-induced bursting activity recorded intracellularly in L1-L2 ventromedial interneurons from wild-type (n = 13 cells) and Kv7.2^Thr274Met/+^ mutant mice (n *=* 18 cells). (**I**) Boxplots quantification of burst parameters. \**P* < 0.05, comparing wild-type *versus* mutant; Mann-Whitney test. Underlying numerical values can be found in the Figure 6 source data.

The above predictions were explicitly tested by means of intracellular recordings of pacemaker cells driven by *I*_NaP_ and triggered by removing the Ca^2+^ from extracellular solution [5, 9, 43]. In this recording condition the burst termination did not involve *I*_*KCa*_, but engaged a K^+^ current as the broad-spectrum K^+^ channel blocker TEA strongly increased the duration of bursts, until ultimately a plateau-like depolarization developed (*P* < 0.05, **S5C and S5D Fig**). This result led us to consider the involvement of a non-inactivating TEA-sensitive K^+^-current such as *I*_M_ [44]. In line with computational predictions, the blockade *I*_M_ by XE991 increased the duration and decreased the frequency of bursts (*P* < 0.05, **Fig 6E and 6G**). Similar results were obtained with linopirdine (**S5E and S5F Fig**). The ability of ICA73 to achieve the converse through the magnification of *I*_M_ was found (*P* < 0.05, **Fig 6F and 6G**). The effects of ICA73 on burst dynamics were reproduced with retigabine (**S5G and S5H Fig**). Interestingly, as predicted by the model, the enhancement of *I*_M_ by ICA73 converted rhythmic bursting to spiking into few cells (**S5I Fig**). Finally, to investigate the contribution of Kv7.2 channels, pacemaker cells were recorded from Kv7.2^Thr274Met/+^ mice in [Ca^2+^]_o_-free saline. Pacemaker properties displayed long-lasting and low-frequency bursts relative to those recorded in Kv7.2^+/+^ littermates (*P* < 0.05, *P* < 0.01, **Fig 6H and 6I**).

Together these data show that *I*_M_, at least mediated by Kv7.2-containing channels, takes a significant part in the timing/intensity control of bursts to regulate dynamics of oscillatory properties in pacemakers.

### M-current mediated by Kv7.2-containing channels controls the locomotor cycle

To investigate the role of *I*_M_ in networkwide rhythmogenesis, we examined its role in the operation of the locomotor rhythm-generating network by using *in vitro* spinal cord preparations from neonatal rats. Rostral lumbar segments (L1/L2) have a powerful rhythmogenic capacity than the caudal ones in neonatal rats [1, 2]. A Vaseline barrier was built at the L2/L3 level to selectively superfuse the two compartments with different drug cocktails (**Fig 7A**). During bath-application of NMA/5-HT in the two sides of the barrier to induce locomotor-like activities, the addition of XE991 (10 µM) in the highly-rhythmogenic L1/L2 compartment to decrease *I*_M_ led to an augmentation of the locomotor cycle period, burst duration and burst amplitude (*P* < 0.05, **Fig 7B**). Also, when preincubated for 30 min before the application of NMA/5-HT, XE991 strongly decreased the latency for the emergence of fictive locomotion (*P* < 0.05, **S6 Fig**). By contrast, augmenting *I*_M_ with ICA73 sped up locomotor cycles and shortened burst duration without apparent effect on burst amplitude (*P* < 0.05, **Fig 7C**). To study the contribution of Kv7.2 channels, fictive locomotion was induced in Kv7.2^Thr274Met/+^ mice. It appeared slower (*P* < 0.05) with longer burst duration (*P* < 0.01) compared to that induced in wild type spinal cords (**Fig 7D**).

**Figure 7:**
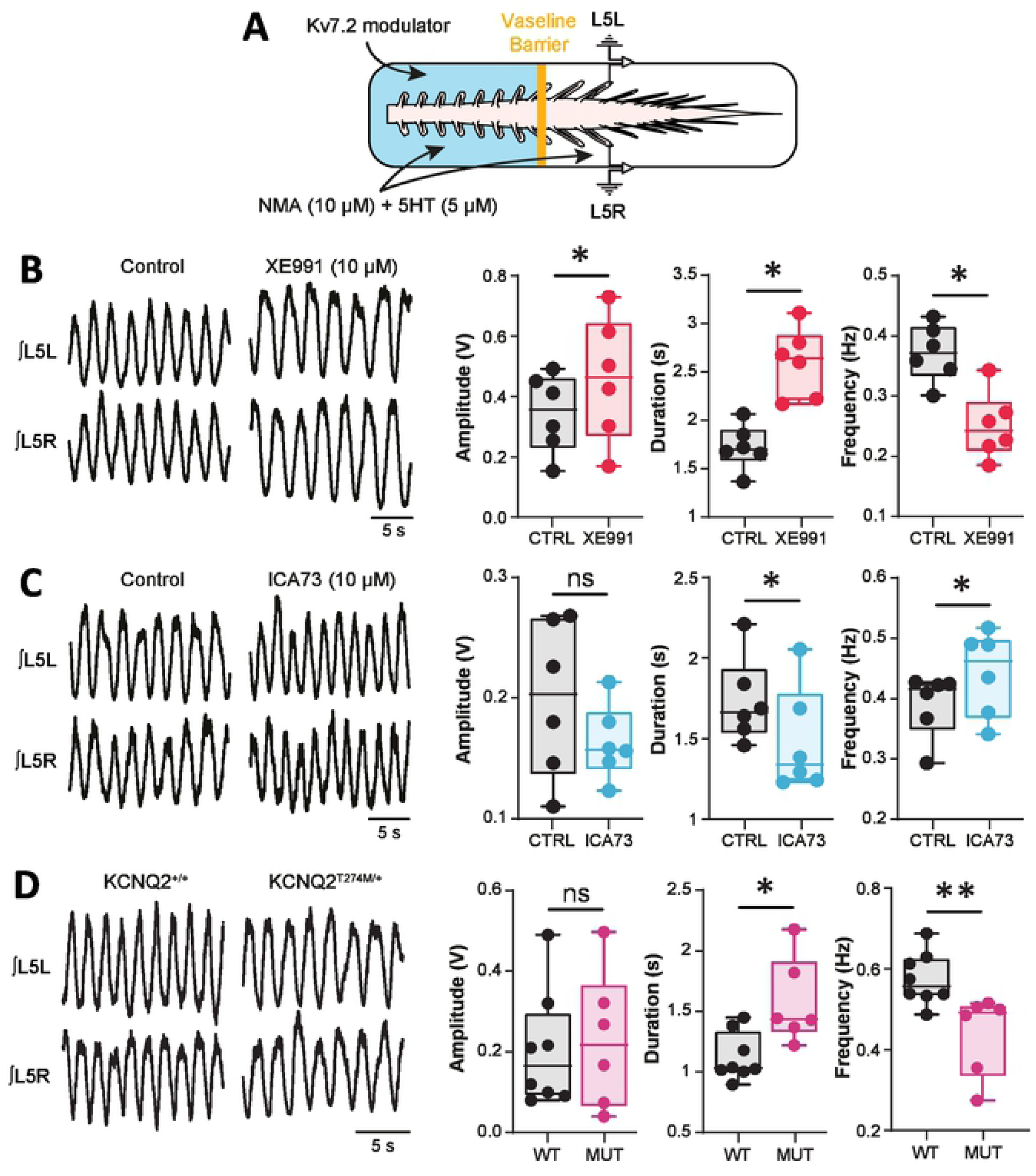
*I*_M_ mediated by Kv7.2 containing channels modulates the locomotor cycle at the level of the CPG. (**A**) Experimental setup. (**B-C**) Ventral-root recordings of NMA/5-HT-induced rhythmic activity generated before and after adding XE991 (10 µM, n = 5 spinal cords) (**B**) or ICA73 (10 µM, n = 6 spinal cords) (**C**) to rostral lumbar segments. On the right, boxplots quantification of locomotor burst parameters. \**P* < 0.05, comparing data before and after the above-mentioned drugs; Wilcoxon paired test. (**D**) Ventral-root recordings of NMA/5-HT-induced rhythmic activity generated in control and Kv7.2^Thr274Met/+^ mice. On the right, boxplots quantification of locomotor burst parameters. \**P* < 0.05, **P < 0.01, comparing wild-type *versus* mutant; Mann-Whitney test. Underlying numerical values can be found in the figure 7 source data.

In some CNS neurons, M-channels have been reported to increase the glutamatergic release by acting at a presynaptic level [45, 46]. Because ionotropic glutamate receptors tune the locomotor network to perform at different speeds [47], we tested the possibility that XE991 and ICA93 affect central glutamatergic synaptic transmission at the CPG level. We analyzed miniature excitatory postsynaptic currents (mEPSCs) recorded from L1–L2 ventromedial interneurons in slice preparations (*P* > 0.05, **S7 Fig**). Neither the amplitudes nor the frequencies of mEPSPCs were affected by Kv7 modulators. Altogether, these results provide new insights into the operation of the CPG with a critical implication of *I*_M_ in the dynamic modulation of the locomotor rhythm.

## Discussion

We provide evidence of the expression of *I*_*M*_ in locomotor-related interneurons such as Hb9 cells and of its critical importance for the operation of the rhythm*-*generating mechanisms. In sum, *I*_*M*_ appears mainly mediated by axonal Kv7.2 channels and acts in opposition to *I*_*NaP*_ by modulating both the emergence and frequency regime of pacemaker cells and thereby regulates the clock function of the spinal locomotor network.

*I*_*M*_ has been previously characterized in sensory and motor neurons of the spinal cord [27, 28, 48, 49] with a significant role in dampening their excitability [26, 28] to prevent both nociception and myokymia [50, 51]. Here, we attempted to identify *I*_*M*_ at the core of the spinal rhythm-generating network for locomotion by recording L1-L2 ventromedial interneurons and Hb9^+^ cells [38, 39]. All of them display a persistent potassium current that shares many characteristics with *I*_*M*_, including voltage dependence, kinetics and pharmacology [52-55]. It has been suggested that native *I*_*M*_ can be mediated by heteromeric assemblies of Kv7.2 and Kv7.3 subunits [53, 56] with an obligatory role of Kv7.2 channels [57]. A co-expression of both Kv7 subunits occurs in spinal dorsal root ganglia and motoneurons Pan [29, 48]. On the other hand, our immunocytochemistry showed that most of neurons within the CPG region are endowed of Kv7.2 subunits but weakly expressed, if any, Kv7.3 subunits. Although we cannot exclude the possibility of some contribution by Kv7.3 channels to *I*_*M*_, the prevalent contribution by Kv7.2 channels is most likely. This is supported by the decrease *I*_*M*_ in CPG cells recorded from KCNQ2^T274M/+^ mutant mice. In sum, our data plead for a dominant contribution of Kv7.2 channels carrying the native *I*_*M*_ in CPG neurons of juvenile rodents, possibly as homomeric form.

In our model, the omission of *I*_*M*_ depolarized the resting membrane potential predicting a contribution of *I*_*M*_ in setting this parameter. Such a role, previously described for *I*_*M*_ in several neuronal systems [58-63], assumes a steady-state activation of the outward current at rest. This is in agreement with the threshold activation of *I*_*M*_ that was found slightly more negative than the resting membrane potential in ventromedial and Hb9 interneurons. On the other hand, the pharmacological inhibition of *I*_*M*_ did not affect their resting membrane potential. This discrepancy likely results from the inefficiency of Kv7 inhibitors to block *I*_*M*_ at membrane potentials close to the resting potential [41, 42]. The *I*_*M*_ did not appreciably alter the threshold or waveform of action potentials probably due to the slow activation kinetics of Kv7 channels [53]. However, *I*_*M*_ dampens neuronal excitability of CPG cells by tuning the threshold current required to fire an action potential. In fact, as originally described in frog sympathetic neurons [64], *I*_*M*_ impacted neuronal excitability of CPG cells primarily by hyperpolarizing their membrane and reducing their input resistance. In sum, *I*_*M*_ appears to be an important player to control both the excitability and the firing frequency at which CPG cells are able to fire.

The present study supports a central role for *I*_*M*_ in shaping the firing pattern of CPG interneurons. In response to a sustained depolarization, a decrease of *I*_*M*_ converts a spiking pattern into a bursting mode in a subset of ventromedial spinal interneurons. Within both cortical and hippocampal pyramidal neurons in mammals, the prevalence of bursting also increases after a pharmacological [46, 59, 61, 62, 65] or a genetic alteration of *I*_*M*_ [57]. Therefore, a basal Kv7.2 channel activity acts as a “brake” in controlling the bursting behavior of CPG cells. This assumption accords with indirect evidence suggesting that the inhibition of a M-like current might regulate pacemaker bursting cells in spinal cord primary cultures [66].

Because *I*_NaP_ functions as the primary mechanism for oscillatory burst generation in CPG interneurons [3, 5, 8, 9], it is conceivable that *I*_*M*_ interacts with *I*_NaP_ to orchestrate bursting behavior. Several pieces of evidence support the existence of a dynamic interplay between *I*_*M*_ and *I*_NaP_. First, our voltage clamp recordings showed a facilitation of *I*_NaP_ once *I*_*M*_ is reduced. Thus, even if the activation of Kv7 channels is too slow to influence the transient sodium current associated with the spike generation (see above), *I*_*M*_ appears fast enough to interact with *I*_NaP_. Second, according to our model that incorporates a heterogeneous distribution of *I*_NaP_ and *I*_*M*_, the principal distinguishing property between bursting pacemaker versus nonpacemaker behaviors was the relative magnitude of *I*_NaP_ to *I*_*M*_; that is, bursting cells displayed a higher *I*_NaP_/*I*_*M*_ ratio compared to nonbursting cells. Third, a co-expression between sodium and Kv7.2 channels was found in interneurons from the CPG region at the AIS, supposed to be the primary source for both *I*_*M*_ and *I*_NaP_ [61, 62, 67, 68]. Altogether, our data indicate that *I*_NaP_ and *I*_*M*_ are ubiquitously expressed in CPG neurons, and that the core biophysical mechanism for oscillatory activities relies on the spatial and temporal dynamic interactions between the two conductances.

In addition to regulate the prevalence of bursting cells in the CPG, *I*_*M*_ predominantly influences bursting dynamics of pacemaker neurons. At the cellular level, the decrease of *I*_*M*_ delays the burst termination suggesting that *I*_*M*_ helps to swing the membrane potential down. Another notable consequence was the increase in the interburst interval even though *I*_*M*_ appears negligible during this period. This paradox implies that *I*_*M*_ indirectly dictates the interburst interval. Since Na^+^ is the main charge carrier for burst generation, we posit that *I*_*M*_ indirectly influences the interburst interval by modulating the activity-dependent post-burst hyperpolarization mediated by Na^+^ /K^+^ pumps in spinal neurons notably within the CPG [21, 22, 69, 70]. A parallel Na^+^-dependent K^+^ current might also participate [71, 72].

Together, these observations support that *I*_*M*_ is part of the burst firing-activated outward current contributing to the burst termination. Rather than exclusively controlled by *I*_*M*_, the burst termination relies on complementary factors acting in concert. Consistent with this, the substantial increase in burst duration induced by TEA, until ultimately reaching a depolarization block, assumes the contribution of voltage-gated K^+^ currents others than *I*_*M*_ in the burst termination process. This observation is consistent with a previous report identifying at least three K^+^ conductances (*I*_A_; *I*_Kdr_; *I*_KCa_) controlling the activity of the bursting in cultured spinal neurons [73]. On the other hand, in our recording conditions performed in [Ca^2+^]_o_-free saline, no obvious role could be attributed to *I*_KCa_. In conclusion, we suggest that the burst repolarization in CPG cells is related to activation of multiple K^+^ currents including *I*_*M*_.

Rhythmic motor systems are characterized by the ability to regulate the cycle frequency of the rhythm. Multiple outward currents contribute to modulate the locomotor rhythm [14, 15, 18, 20], but the involvement of *I*_*M*_ in the locomotor function has never been investigated. The specific modulation of the fictive locomotor rhythm following the selective application of Kv7-modulators over the CPG demonstrates that *I*_*M*_ represents a new ionic component by which the speed of locomotion can be tuned. The effects of *I*_*M*_ on the controllability of the locomotor rhythm *in vivo* being reminiscent of those observed on fictive locomotion, lead us to conclude that *I*_*M*_ provides not only a mechanism for rhythmic bursting of individual pacemaker cells, but also for rhythmic activity at the network level to adapt speed of movements as circumstances demand. This investigation is the first direct evidence for the concept that *I*_*M*_ plays a key role in controlling rhythmogenesis in the spinal locomotor network. *I*_*M*_ is a well-established target for a range of modulators [74-77], and thus may offer a powerful mean to regulate the rhythmicity of the spinal locomotor network. Regarding the initial characterization of *I*_*M*_ through its suppression by muscarinic receptor activation [64, 78] one obvious candidate is acetylcholine. The cholinergic system of the spinal cord commonly produces locomotion in a slow speed range with a significant contribution of muscarinic receptors [79-82]. Because the inhibition of *I*_*M*_ slowed down the locomotor rhythm, the muscarinic inhibition of *I*_*M*_ at the CPG level might play a key role in the dynamic reconfiguration of the locomotor network by changing the relative number of bursting pacemaker cells, and therefore the degree of their engagement in network activity [83]. In line with this notion, cholinergic cells are active during locomotion [84, 85] and a significant proportion of CPG neurons are responsive to acetylcholine in the form of intrinsic membrane potential oscillations [81, 86]. Monoamines such as serotonin also shape spinal motor patterns in mammals particularly by lengthening their locomotor rhythm [87]. Considering the ability of the monaminergic system to interact with Kv7 channels [88], and of serotonin to promote burst firing through a decrease of *I*_*M*_ [89], it is conceivable that neuromodulation of *I*_*M*_ by monoamines dynamically reconfigures the firing pattern of locomotor CPG interneurons.

Aside neuromodulation, we indicated that the propensity of a CPG neuron to burst also depends on the ionic composition of the milieu in which it is embedded [3-5, 9]. As a consequence of activity-dependent changes in extracellular calcium ([Ca^2+^]_o_) and potassium ([K^+^]_o_) concentrations during locomotion, a large number of CPG interneurons are converted from regular-spiking into bursting through a concomitant upregulation of *I*_NaP_ and reduction of K^+^ currents [9]. Thus, by modulating *I*_NaP_ and *I*_*M*_, respective changes in [Ca^2+^]_o_ and [K^+^]_o_ may represent a fast and powerful mechanism to regulate bursting pacemaker cells and thereby the operation of the locomotor CPG. Overall, this study provides new insights into the operation of the locomotor network whereby *I*_M_ and *I*_NaP_ represent a functional set of subthreshold currents that endow the locomotor CPG with rhythmogenic properties, with a behavioral role of *I*_M_ in controlling the speed of locomotion.

## Acknowledgments

We are grateful the lab members for their critical reading of the manuscript and Anne Duhoux for animal care.

## Author contributions

J.V. performed and analyzed most of *in vitro* and *in vivo* experiments. J.V. performed and analyzed *in silico* experiments. C.B. performed and analyzed immunohistochemistry experiments with J.V. L.V. provided the genetic model of I_M_ dysfunction. F.B. designed and supervised the whole project with J.P.R.. F.B. wrote the manuscript.

## Material and Methods

### Ethics statement

We made all efforts to minimize animal suffering and the number of animals used. All animal care and use conformed to the French regulations (Décret 2010-118) and were approved by the local ethics committee (Comité d’Ethique en Neurosciences INT-Marseille, CE Nb A1301404, authorization Nb 2018110819197361). Experiments were performed on Wistar rats, Hb9:eGFP mice and heterozygous KCNQ2^T274M/+^ mutant mice.

### The Kcnq2^Thr274Met/+^ mouse model

It was generated by homologous recombination in embryonic stem (ES) cells using a targeting vector containing regions homologous to the genomic *Kcnq2* sequences and the p.(Thr274Met) variant, a recurrent pathogenic variant identified in several patients suffering from developmental and epileptic encephalopathy. Correctly targeted 129Sv ES cell clones were injected into C57Bl/6N blastocysts implanted in pseudo-pregnant females. Chimerism rate was assessed in the progeny by coat colour markers comparison and the mice were bred with 129sv Cre-deleter mice to excise the Neomycin selection cassette and to generate *Kcnq2*^Thr274Met/+^ mice. Genotyping was performed using genomic DNA prepared from ear punch biopsies with the Direct DNA (Tail) (Viagen Biotech, USA). The *Kcnq2*^Thr274Met/+^ animals were maintained and studied on the 129Sv genetic background. Wildtype and heterozygous knock-in animals express the same amount of Kcnq2 transcript and the two alleles are equally expressed. The characterization of the *Kcnq2*^Thr274Met/+^ mouse reveals that it faithfully reproduces what is expected based on the human phenotype: no gross morphological brain alterations, no neurosensory alterations before the onset of seizures occurring at P20 followed by a high rate of unexpected death in epilepsy and important cognitive difficulties [40].

### Assessment of locomotor behaviors

Animals were tested in juvenile animals (P15– P21) when a mature pattern of locomotion occurred [90]. The CatWalkXT (v9.1, Noldus Information Technology, Netherlands) was employed to measure walking performance. Each animal walked freely through a corridor on a glass walkway illuminated with beams of light from below. A successful walking trial was defined as having the animal walk at a steady speed (no stopping, rearing, or grooming), and three to five successful trials were collected per animal. Experimental sessions typically lasted for 5 to 10 min. The footprints were recorded using a camera positioned below the walkway, and footprint classification was manually corrected to ensure accurate readings. The paw print parameters were then analyzed using the CatWalk software (see data analysis).

#### *In vitro* models

Details of the *in vitro* preparations have been previously described vinay [91, 92] and are only summarized here. Experiments were performed from newborn rats or mice (1-to 5-day-old). *<u>For the whole spinal cord preparation</u>*, the spinal cord was transected at T10, isolated and transferred to the recording chamber perfused with oxygenated artificial cerebrospinal fluid (aCSF). For rats, the aCSF was composed of (in mM): 120 NaCl, 4 KCl, 1.25 NaH_2_PO_4_, 1.3 MgSO_4_, 1.2 CaCl_2_, 25 NaHCO_3_, 20 D-glucose, pH 7.4 (25–26°C). For mice, the aCSF was composed of (in mM): 128 NaCl, 4 KCl, 0.5 NaH_2_PO_4_, 1 MgSO_4_, 1.5 CaCl_2_, 21 NaHCO_3_, 30 D-Glucose, pH 7.4 (25–26°C). *<u>For the slice preparation</u>*, the lumbar spinal cords was isolated in ice-cold (<4°C) aCSF with the following composition (in mM): 232 sucrose, 3 KCl, 1.25 KH_2_PO_4_, 4 MgSO_4_, 0.2 CaCl_2_, 26 NaHCO_3_, 25 D-glucose, pH 7.4. The lumbar spinal cord was then introduced into a 1% agar solution, quickly cooled, mounted in a vibrating microtome (Leica VT1000S) and sliced (350 µm) through lumbar segments. Slices were immediately transferred into the holding chamber filled with aCSF composed of (in mM): 120 NaCl, 3 KCl, 1.25 NaH_2_PO_4_, 1.3 MgSO_4_, 1.2 CaCl_2_, 25 NaHCO_3_, 20 D-glucose, pH 7.4 (32-34°C). Following a 1 h resting period, individual slices were transferred to a recording chamber that was continuously perfused with the same medium heated to ∼27°C. Slices were visualized with epifluorescence and infrared differential interference contrast (IR-DIC) microscopy using a Nikon Eclipse E600FN upright microscope coupled with a 40 X water immersion lens. The image was enhanced with an infrared-sensitive CCD camera and displayed on a video monitor. The temperature regulation was provided by the CL-100 bipolar temperature controller (Warner Instruments).

#### *In vitro* recordings

*<u>For the whole spinal cord preparation</u>*, motor outputs were recorded from lumbar ventral roots (left / right L5) by means of glass suction electrodes connected to an AC-coupled amplifier. The ventral root recordings were amplified (×2,000), high-pass filtered at 70 Hz, low-pass filtered at 3 kHz, and sampled at 10 kHz. Custom-built amplifiers enabled simultaneous online rectification and integration (100 ms time constant) of raw signals. Locomotor-like activity was induced by a bath application of N-methyl-DL aspartate (NMA, 10 μM) and 5-hydroxytryptamine (5-HT, 5 μM). In some experiments, a Vaseline barrier was built at the L_2_/L_3_ level to superfuse the locomotor network located in the rostral lumbar cord independently from the more caudally located motoneurons. *<u>For the slice preparation</u>*, whole-cell patch-clamp recordings were performed from L1-L2 interneurons using a Multiclamp 700B amplifier (Molecular Devices). Interneurons located in the medial lamina VIII and adjacent to the central canal, a region proposed to contain a large part of the rhythm-generating locomotor network [2], were selected. In transgenic Hb9:eGFP transgenic mice, only GFP-positive interneurons compatible with the previously described electrophysiological profile of Hb9 interneurons i.e. high input resistance, strong post-inhibitory rebound and absence of sag were considered [93]. Motoneurons were visually identified as the largest cells located in layer IX. Patch electrodes (2-4 MΩ) were pulled from borosilicate glass capillaries (1.5 mm OD, 1.12 mm ID; World Precision Instruments) on a Sutter P-97 puller (Sutter Instruments Company) and filled with intracellular solution containing (in mM): 140 K^+^-gluconate, 5 NaCl, 2 MgCl_2_, 10 HEPES, 0.5 EGTA, 2 ATP, 0.4 GTP, pH 7.3 (280 to 290 mOsm). Pipette and neuronal capacitive currents were canceled and, after breakthrough, the series resistance was compensated and monitored. Recordings were digitized on-line and filtered at 10 kHz (Digidata 1322A, Molecular Devices). The main characterization of *I*_M_ was accomplished by holding the membrane potential at a relatively depolarized potential (*V*_H_, -10 mV) to activate KCNQ channels and to inactivate many of the other K^+^ channels notably Kv1.2 channels [19]. The membrane potential was then stepped down to more hyperpolarized potentials to deactivate the KCNQ channels giving rise to slow inward current relaxation. Stepping back to -10 mV led to the reactivation of the KCNQ channels to produce slow inward relaxations. All experiments were designed to gather data within a stable period (i.e. at least 5 min after establishing whole-cell access). Action potential-independent miniature excitatory (mEPSCs) were recorded in the presence of TTX (1 µM) at a holding potential of -70 mV. NMDA and non-NMDA receptor-mediated mEPSCs were recorded with a K^+^-gluconate based intracellular solution (see above) and pharmacologically isolated with a combination of biccuculine (20 µM) and strychnine (1 µM) to fast GABAergic and glycinergic synapses, respectively.

#### Immunohistochemistry

Spinal cords were immersion-fixed for 1 h in 0.25% PFA, washed in PBS and cryoprotected overnight at 4 °C in 20% sucrose in PBS. Lumbar spinal cords (L1-L2) were then frozen in OCT medium (Tissue Tec), cryosectioned (20 µm) and processed for immunohistochemistry. Slices were then (i) rehydrated in PBS at room temperature (15 min), (ii) permeated with 1% Bovin Serum Albumin (BSA), 2% Natural Goat Serum (NGS) and 0.2% Triton x-100 (1 h), (iii) incubated overnight at 4 °C in the following affinity-purified rabbit K_V_7.2 (residues 1–70; 1:1000, Thermo Scientific), K_V_7.3 (residues 668–686; 1:400, Alomone) specific polyclonal antibody, mouse pan Na_v_ (residues 932–1043; 1:1000, Sigma) specific monoclonal antibody (iv) washed in PBS (3×5 min), (v) incubated with fluorescent-conjugated secondary antibodies [Alexa 488-or 546-conjugated mouse-or rabbit-specific antibodies (1:800 and 1:400; Lifetechnologies Carlsbad CA USA) used for visualization of the mouse monoclonal or rabbit polyclonal antibodies, respectively] in a solution containing 1% BSA and 2% NGS (1.5 h), (vi) washed in PBS 3×5 min, (vii) coverslipped with a gelatinous aqueous medium. In control experiments, the primary antiserum replaced with negative control mouse or rabbit immunoglobulin fraction during the staining protocol. Sections were scanned using a laser scanning confocal microscope (Zeiss LSM700) in stacks of 0.3-1 μm-thick optical sections at × 63 and/or × 20 magnification respectively and processed with the Zen 12.0 software (Zeiss). Each optical section resulted from two scanning averages. Each figure corresponds to a projection image from a stack of optical sections.

#### Cell model

The model of the pacemaker neuron is a typical somatic single-compartment model developed in the Hodgkin-Huxley style and was based on our previous study on Hb9 cells [9]. The neuronal membrane potential *V* was dynamically defined by a set of membrane ionic currents. The current balance equation is :

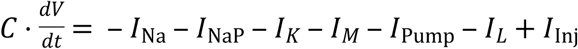

where *C* is neuronal membrane capacitance (pF) and *t* is time (ms). The modeled neuron included the following ionic currents: transient sodium current (*I*_Na_ with the maximal conductance 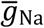), the persistent sodium current (*I*_NaP_ with the maximal conductance 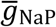), the non-inactivating M-current (*I*_M_ with the maximal conductance 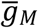), the delayed rectifier potassium current (*I*_K_ with the maximal conductance 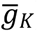), the sodium-dependent pump (*I*_Pump_), leakage (*I*_L_ with the conductance *g*_L_), and depolarizing injected (*I*_Inj_) currents. These currents, except for *I*_Pump_ and *I*_Inj_ are described as follows:

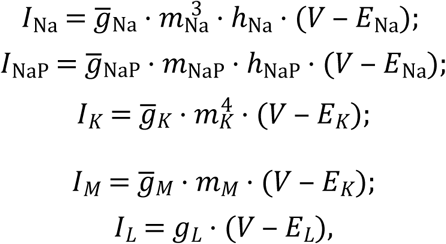

*E*_Na_, *E*_K_, and *E*_L_ are reversal potentials of the corresponding channels (in mV):

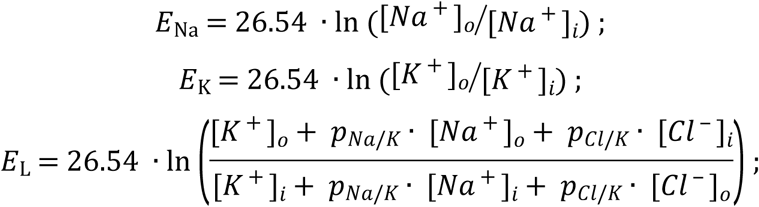

[Na^+^], [K^+^], and [Cl^-^] represent concentrations of sodium, potassium, and chloride ions, respectively. Indexes “o” and “i” define the concentrations of these ions outside and inside a cell, respectively; [Na^+^]_o_= 145 mM, [K^+^]_o_= 3 mM, [K^+^]_i_= 140 mM, [Cl^-^]_i_= 8 mM, [Cl^-^]_0_= 130 mM, [Na^+^]_i_ is considered variable. Parameters *p*_Na/K_ and *p*_Cl/K_ represent the relative permeability of sodium and chloride ions (with respect to potassium ions); *p*_Na/K_ = 0.03, *p*_Cl/K_ = 0.1.

In our model, the intracellular sodium concentration [Na^+^]_i_ is accumulating due to *I*_Na_ and *I*_NaP_ currents and is pumped out by *I*_Pump_. The contributions of these components are defined by the corresponding coefficients (*α*_Na_, *α*_NaP_, and *α*_Pump_ respectively):

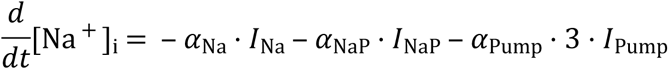

The pump current is described as: *I*_Pump_ = *R*_Pump_ · (*ϕ*([Na ^+^]_*i*_) − *ϕ*([Na ^+^]_ibase_) where 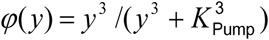 is the base intracellular sodium concentration, *R*_Pump_ and *K*_Pump_ are the *I*_Pump_ parameters. The following parameters were used for sodium dynamics and pump current: *α*_Na_ = *α*_NaP_ = *α*_Pump_ = 10^−5^ mM/fC ; *R*_Pump_ = 60 pA, [Na+]_ibase_= 15 mM, *K*_Pump_ = 18 mM.

The dynamics of activation (*m*) or inactivation (*h*) variables for the above sodium and potassium channels are generally described by the differential equation: *dx*/*dt* = (*x*_∞_ − *x*)/*τ*_*x*_, *x* = {*m,h*}, where *x*_∞_ is the voltage-dependent steady state value and *τ*_*x*_ is the voltage-dependent time constant of the variable *x*, which are described in the following form:

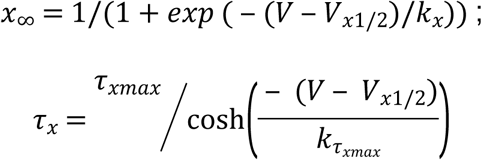

*V*_x1/2_ and *k*_x_ are the half-activation voltage and the slope for variable *x, τ*_*x*max_ is the maximum value of its time constant and *k*_τ_ defines the slope of this time constant. The activation of sodium currents (*I*_Na_ and *I*_NaP_) is considered instant, i.e. *τ*_*m*Na_= *τ*_*m*NaP_ =0, thus *m*_Na_, and *m*_NaP_ are considered equal to their steady state values.

To reproduce our experimental finding, the half-activation voltage for *I*_NaP_, *V*_*m*NaP1/2_, is made depended on the outside calcium concentration:

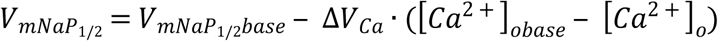

where *V*_mNaP1/2base_ corresponds to the half-activation voltage at [Ca^2+^]_o_=[Ca^2+^]_obase_ =1.2 mM and Δ*V*_Ca_ = 1 mV/mM defines a shift in *V*_mNaP1/2_ occurring with changes in [Ca^2+^]_o_.

In our simulations, we considered either a single neuron or a population of 50 uncoupled neurons. To provide a necessary heterogeneity in properties of neurons, we Gaussian-distributed the base values V_1/2_ for *I*_M_ and *I*_NaP_ derived from our recordings. An additional heterogeneity was set by normal distribution of all conductances around appropriate base values (see Table below). Parameters for *I*_Na_, *I*_K_, and inactivation of *I*_NaP_ were taken from previous modeling studies with some modifications.

| Current | Parameters |
| --- | --- |
| Fast Na<sup>+</sup> | $g_{Na} = 110 \pm 11 \text{ nS}$ |
| | *Act.* $V_{mNa1/2} = -43.8 \text{ mV}$; $k_{mNa} = 6 \text{ mV}$ |
| | *Inact.* $V_{hNa1/2} = -67.5 \text{ mV}$; $k_{hNa} = -10.8 \text{ mV}$; $\tau_{hNa_{max}} = 35.2 \text{ ms}$; $k_{ThNa} = 12.8 \text{ mV}$ |
| Persistent Na<sup>+</sup> | $g_{NaP} = 0.4 \pm 0.1 \text{ nS}$ |
| | *Act.* $V_{mNaP1/2_{base}} = -52 \pm 1.5 \text{ mV}$; $k_{mNaP} = 3 \text{ mV}$ |
| | *Inact.* $V_{hNaP1/2} = -59 \text{ mV}$; $k_{hNaP} = -5.2 \text{ mV}$; $\tau_{hNaP_{max}} = 20000 \text{ ms}$; $k_{ThNaP} = 10.4 \text{ mV}$ |
| K<sup>+</sup> delayed rectifier | $g_K = 60 \pm 6 \text{ nS}$ |
| | *Act.* $V_{mK1/2} = -34.5 \text{ mV}$; $k_{mK} = 5 \text{ mV}$; $\tau_{mK_{max}} = 4 \text{ mV}$; $k_{tmK} = 30 \text{ mV}$ |
| M current | $g_M = 0.8 \pm 0.1 \text{ nS}$ |
| | *Act.* $V_{mM1/2} = -44 \pm 1.5 \text{ mV}$; $k_{mM} = 4.3 \text{ mV}$; $\tau_{mM_{max}} = 7 \text{ mV}$; $k_{tmM} = 25 \text{ mV}$ |
| Other | $C = 21.5 \text{ pF}$; $g_L = 1 \pm 0.1 \text{ nS}$ |

Simulations were performed using MATLAB R2019b (The MathWorks, Natick, MA). Differential equations were solved using a variable order multistep differential equation solver ode15s available in MATLAB. In each simulation, a settling period of 30 s was allowed before data were collected. For population, each simulation was repeated 10 times, and demonstrated qualitatively similar behavior for particular values of parameters within the standard deviations.

### Data analysis

#### Quantitative gait analyses

The CatWalk XT software (v9.1, Noldus Information Technology, Netherlands) was used to measure a broad number of spatial and temporal gait parameters in several categories. These include i) dynamic parameters related to individual paw prints, such as duration of the step cycle with the respective duration of the swing and stance phases; ii) parameters related to the position of paw prints with respect to each other, for example the stridverneuile length (distance between two consecutive placement of the same paw) and the base of support (the width between both the front and hind paws); iii) parameters related to time-based relationships between paw pairs, as well as step patterns. These parameters were initially calculated for each run and for each paw, then averaged over the runs, and finally the values for the front and hind paws were averaged. To determine which parameters were affected data were normalized against basal values, which were settled as 100% in each parameter.

#### Electrophysiological data analyses

The Clampfit 10.7 software (Molecular devices) was used for analyzing electrophysiological data. Alternating activity between right/left L5 recordings was taken to be indicative of fictive locomotion. To characterize locomotor burst parameters, raw extracellular recordings from ventral roots were rectified, integrated and resampled. Peak amplitude of locomotor burst was measured and the cycle period was calculated by measuring the time between the first two peaks of the auto-correlogram. The coupling between right/left L5 was estimated by measuring the correlation coefficient of the cross-correlogram at zero phase lag. The onset of a locomotor-like activity was determined when a clear rhythmic alternating activity was observed. Several basic criteria were set to ensure optimum quality of intracellular recordings. Only cells exhibiting a stable resting, holding membrane potential, access resistance (less than 20% variation) and an action potential amplitude larger than 40 mV were considered. All reported membrane potentials were corrected for liquid junction potentials. We determined input resistance by the slope of linear fits to small (<5mV) voltage responses evoked by positive and negative current injections. Firing properties were measured from depolarizing current pulses of varying amplitudes. The rheobase was defined as the minimum step current intensity required to induce an action potential from the membrane potential held at -60mV. Single spike analysis was performed on the first spike elicited near the rheobase. Peak spike amplitude was measured from the threshold potential, and spike duration was defined as the time to fall to half-maximum peak. The instantaneous discharge frequency was determined as the inverse of interspike interval and plotted as a function of time. For a direct comparison of firing properties before and during the application of the drug, a bias current could be used to maintain the membrane potential at the holding potential fixed in the control condition. Voltage dependence and kinetics of *I*_M_ were analyzed from data normalized to the maximal current. The I/V curves were fitted with a Boltzmann function. *I*_M_ amplitude was measured from deactivation relaxation at -50mV. mEPSCs were detected and analyzed using the MiniAnalysis Program (Synaptosoft, NJ, USA). Events were detected by setting the threshold value for detection at 3 times the level of the root mean square noise (∼3-4pA meaning detection threshold at ∼8-12pA). The average values of mPSCs amplitude and frequency during the control period and 30 min after the drug application, were calculated over a 5-min time window.

#### Immunohistochemistry analysis

Neurons with AISs of obvious soma origin were imaged. Image stacks were converted into single maximum intensity z-axis projections and imported into Matlab (Mathworks) for analysis using a previously published MATLAB code [94] downloaded from the Grubb Lab (ais_z3.m from http://grubblab.org/resources/). Measurements were performed on initial segments from interneurons located in the ventromedial part of upper lumbar segments (L1-L2) near the central canal and from motoneurons identified as the biggest cells located in the ventral horn. Initial segments were identified as linear structures labeled by pan Na_v_-specific antibodies, and for which the beginning and the end of the structure could be clearly determined, excluding nodes of Ranvier. We drew a line profile starting at the soma that extended down the axon, through and past the AIS. For quantifications of the start and end position of immunolabelings along the axonal process from the soma, axonal profiles were smoothed using a ∼5 μm sliding mean and normalised to the maximum smoothed fluorescence. AIS start and end positions were identified as the points where fluorescence intensities increased above and dropped below 33% of the maximum axonal fluorescence intensity, respectively, in line with previous reports [94].

### Drug list and solutions

Normal aCSF was used in most cases for *in vitro* electrophysiological recordings. Ca^2+^-free solution was made by removing Ca^2+^ chloride from the recording solution and replacing it with an equimolar concentration of magnesium chloride. All solutions were oxygenated with 95% O_2_/5% CO_2_. For whole-cell voltage-clamp recordings of *I*_M_, the aCSF was supplemented with 1 µM tetrodotoxin (TTX). All salt compounds, tetraethylammonium (TEA) acid kynurenic; strychnine; biccuculine; riluzole; N-methyl-DL-aspartic acid (NMA) and 5-hydroxytryptamine creatinine sulfate (5-HT) were obtained from Sigma-Aldrich. Other drugs, including 10,10-bis(4-pyridinylmethyl)-9(10H)-anthracenone dihydrochloride (XE-991), 1,3-Dihydro-1-phenyl-3,3-*bis*(4-pyridinylmethyl)-2*H*-indol-2-one dihydrochloride (Linopirdine), *N*-(2-Chloro-5-pyrimidinyl)-3,4-difluorobenzamide (ICA-069673 or ICA73), Ethyl [2-amino-4-[[(4-fluorophenyl)methyl]amino]phenyl] carbamate (Retigabine), 2-amino-6-trifluoromethoxybenzothia-zole hydrochloride (riluzole), and TTX were obtained from Tocris Bioscience. Riluzole, Retigabine and ICA73 were dissolved in dimethylsulphoxide (DMSO) and added to the aCSF (final concentration of DMSO: 0.05%). The other drugs were dissolved in water and added to the aCSF. Control experiments showed no effects of the vehicle (data not shown).

### Treatment design

Juvenile rats (15-to 21-day-old) were randomly treated with a single dose of linopirdine, retigabine, XE-991 or ICA73 or its vehicle. The dose of drugs refers to previous reports [30-32]. All the drugs and their vehicle were administered intraperitoneally. The behavioral test was performed before drug treatment and 30 min after the i.p. injection of drug or vehicle.

### Statistics

No statistical method was used to predetermine sample size. Group measurements were expressed as means ± s.e.m. We used Mann-Whitney test or Wilcoxon matched pairs test to compare two groups. For all statistical analyses, the data met the assumptions of the test and the variance between the statistically compared groups was similar. The level of significance was set at *P* < 0.05. Statistical analyses were performed using Prism 5.0 software (Graphpad). Boxplots in figures show the distribution extremes (horizontal bars), 25th and 75th percentiles (box height), and median (center bar).

## Supporting Information

**S1 Fig: *I***_**M**_ **does not affect inter-limb coordination and gait.** (**A-D**) Normalized changes of the stride length (**A**), the base of support (**B**), the number of normal step sequence patterns (**C**) and regularity index of paw placements (**D**) during CatWalk locomotion of juvenile rats before and 30 min after acute i.p. administration of DMSO (*grey, n* = 6 rats), retigabine (5 mg/kg, *green, n* = 7 rats), linopirdine (3 mg/kg, *yellow, n* = 6 rats), ICA73 (5 mg/kg, *blue, n* = 7 rats) or XE991 (5 mg/kg, *red, n* = 6 rats). Dashed lines with grey shading indicate the 95% confidence intervals of control values.n.s. *P* > 0.05, comparing data collected before and after drug administration; Wilcoxon paired test. Underlying numerical values can be found in the S1 Fig source data.

**S2 Fig: Lumbar motoneurons express Kv7.2-containing channels.** (**A-F**) Immunostaining of lumbar (L1-L2) motoneurons from juvenile rats (n = 3 rats) against Kv7.2 (**A**, n = 85 cells) or Kv7.3 (**D**, n = 104 cells) along the AIS labeled by the Pan-Nav antibody (**B**,**E**). Kv7.2 and Pan-Nav are merged in (**C**), and Kv7.3 and Pan-Nav are merged in (**F**). Asterisks indicate the nucleus position and arrowheads the AIS. Scale bars = 20 µm. (**G**) Group means quantification of the proportion of Pan-Nav positive motoneurons expressing Kv7.2 or Kv7.3 channels. (**H**) Group means quantification of the start and end positions of Pan-Nav, Kv7.2 and Kv7.3 immunolabelings along the axonal process from the soma (n = 10 cells). \**P* < 0.05, **P < 0.01, comparing start or end positions between groups; Mann-Whitney test. Data are mean ± SEM. Underlying numerical values can be found in the S2 Fig source data.

**S3 Fig: The *I***_**M**_ **enhancer retigabine replicates the effect of ICA73 on electroresponsive properties of L1-L2 ventromedial interneurons.** (**A**) Representative deactivation of *I*_M_. (**B**), Boltzmann fitted current-voltage relationships of *I*_M_. (**C-F**) Boxplots quantification of the amplitude (**C**), the holding current (**D**), threshold (**E**) and V_1/2_ (**F**) of *I*_M_ recorded in interneurons (n = 7 cells) before (black) and after bath-applying retigabine (100 nM, green). \**P* < 0.05; Wilcoxon paired test. (**G, H**) Typical spiking activity of ventromedial interneurons (L1-L2) to a near threshold depolarizing pulse (**G**) with the respective frequency-current relationship (**H**) before (black) and after bath-applying retigabine (100 nM, green). The continuous line is the best-fitting linear regression. \*\*\**P* < 0.001, comparison of the fits. Underlying numerical values can be found in the S3 Fig source data.

**S4 Fig: The *I***_**M**_**-blocker XE991 antagonizes the effect of ICA73 on electroresponsive properties of L1-L2 ventromedial interneurons.** (**A**) Membrane potential changes in response to bath application of ICA73 (10 µM) before (*light blue*) and after (*dark blue*) pretreatment with XE991 (10 µM). (**B**,**C**) Representative spiking activity to a near threshold depolarizing step (**B**) and frequency-current relationship (**C**) recorded under XE991 before (red) and after bath-applying ICA73 (n = 5 cells, dark blue). Continuous lines are the best-fitting linear regression. n.s *P* > 0.05 comparison of the fits. (**D**) Boxplots quantification of the rheobase. n.s *P* > 0.05; Wilcoxon paired test. Underlying numerical values can be found in the S4 Fig source data.

**S5 Fig: *I***_**M**_ **controls burst dynamics. (A)** Firing behavior from a bursting pacemaker neuron model in response to two different values of *g*_M_. (**B**) Dependence of the percentage of bursting cells on *g*_M_ in the heterogeneous population model of 50 neurons. In **A** and **B** V_1/2_ *I*_NaP_ = -54mV. (**C**,**E**,**G**,**I**) Free-[Ca^2+^]_o_ saline-induced bursting activity recorded intracellularly in L1-L2 ventromedial interneurons before and after TEA (10mM, n = 7 cells) (**D**), linopirdine (10 µM, n = 10 cells) (**F**), retigabine (100nM, n = 6 cells) (**H**), or ICA73 (10 µM, n = 9 cells) (**I**). (**D**,**F**,**H**) Boxplots quantification of the duration and frequency of bursts. \**P* < 0.05, **P < 0.01, comparing data before and after the above-mentioned drugs; Wilcoxon paired test. Underlying numerical values can be found in the S5 Fig source data.

**S6 Fig: Downregulation of *I***_**M**_ **facilitates the emergence of locomotor-like activity.** (**A, B**) Ventral-root recordings of NMA/5-HT-induced rhythmic activity generated without (n = 9 spinal cords) (**A**) and with (n = 6 spinal cords) a 45 min preincubation of XE991 (10 µM) (**B**). Broken-line boxes indicates the part of the recordings that is enlarged in insets to visualize the onset of locomotor-like activity (rhythmic alternating activity). (**C**) Boxplots quantification of the delay between the start of the bath-application of NMA/5-HT and the onset of the fictive locomotion. \**P* < 0.05; Mann-Whitney test. Underlying numerical values can be found in the S6 Fig source data.

**S7 Fig: *I***_**M**_ **has no effect on glutamatergic synaptic transmission.** (**A**) Representative current traces of continuously recorded miniature excitatory postsynaptic currents (mEPSCs) in a ventromedial interneuron voltage clamped at -60 mV. mEPSCs recorded before and 30 min after adding XE991 (10 µM, n = 6 cells) or ICA73 (10 µM, n = 6 cells) were pharmacologically isolated in the presence of TTX (1 µM), strychnine (1 µM), and bicuculline (20 µM). (**B**) Averaged mEPSCs recorded before (black) and 30 min after adding XE991 (red) or ICA73 (blue). (**C**) Boxplots quantification of the mean frequency and amplitude of mEPSCs before and 30 min after XE991 or ICA73 was bath-applied. n.s. *P* > 0.05, comparing data collected before and after bath-applying the above-mentioned drug; Wilcoxon paired test. Underlying numerical values can be found in the S7 Fig source data.

